# BURRITO: An interactive multi-omic tool for visualizing taxa-function relationships in microbiome data

**DOI:** 10.1101/217315

**Authors:** Colin P McNally, Alexander Eng, Cecilia Noecker, William C Gagne-Maynard, Elhanan Borenstein

**Affiliations:** Department of Genome Sciences, University of Washington, Seattle, Washington, USA; Institute for Health Metrics and Evaluation, Seattle, Washington, USA; Department of Computer Science and Engineering, University of Washington, Seattle, Washington, USA; Santa Fe Institute, Santa Fe, New Mexico, USA

**Author notes:** These authors contributed equally to this work (Colin P McNally, Alexander Eng, Cecilia Noecker).

**Keywords:** Microbiome, Metagenomics, Data visualization, Taxonomy, Function, Web interface

## Abstract

The abundance of both taxonomic groups and gene categories in microbiome samples can now be easily assayed via various sequencing technologies, and visualized using a variety of software tools. However, the assemblage of taxa in the microbiome and its gene content are clearly linked, and tools for visualizing the relationship between these two facets of microbiome composition and for facilitating exploratory analysis of their co-variation are lacking. Here we introduce *BURRITO*, a web tool for interactive visualization of microbiome multi-omic data with paired taxonomic and functional information. BURRITO simultaneously visualizes the taxonomic and functional compositions of multiple samples and dynamically highlights relationships between taxa and functions to capture the underlying structure of these data. Users can browse for taxa and functions of interest and interactively explore the share of each function attributed to each taxon across samples. BURRITO supports multiple input formats for taxonomic and metagenomic data, allows adjustment of data granularity, and can export generated visualizations as static publication-ready formatted figures. In this paper, we describe the functionality of BURRITO, and provide illustrative examples of its utility for visualizing various trends in the relationship between the composition of taxa and functions in complex microbiomes.

## Background

Microbial communities are complex ecosystems with important impacts on human health and on the environment. High-throughput DNA sequencing has enabled comprehensive profiling of these communities in terms of their composition and structure. Traditionally, microbial ecology studies resort to one of two primary approaches for profiling the composition of a given community, focusing either on its taxonomic composition (e.g., using targeted 16S rRNA gene sequencing or a marker-gene based approach) or on its functional composition (e.g., using metagenomic shotgun sequencing and assessing the abundance of various gene families) (Figure 1A). Obtained taxonomic or functional profiles are then often visualized as simple stacked bar or area plots of relative abundances, or via specialized data visualization tools designed for exploring these data. For example, Explicet (Robertson et al., 2013) and Krona (Ondov et al., 2011) aid analysis by displaying abundance data while simultaneously presenting the hierarchical relationships between entities. Other tools have gone beyond relative abundance visualization to enable exploration of specific, comparative aspects of microbiome data. EMPeror (Vázquez-Baeza et al., 2013), for example, allows users to generate 3D principal coordinate analysis plots to visualize clustering of, or variation in, taxonomic compositions coupled with trends in any associated metadata. Other tools, including Community-Analyzer (Kuntal et al., 2013) and MetaCoMET (Wang et al., 2016), provide access to multiple types of visualizations of the same microbiome data, each highlighting different aspects of community structure or between-sample relationships.

**Figure 1.**
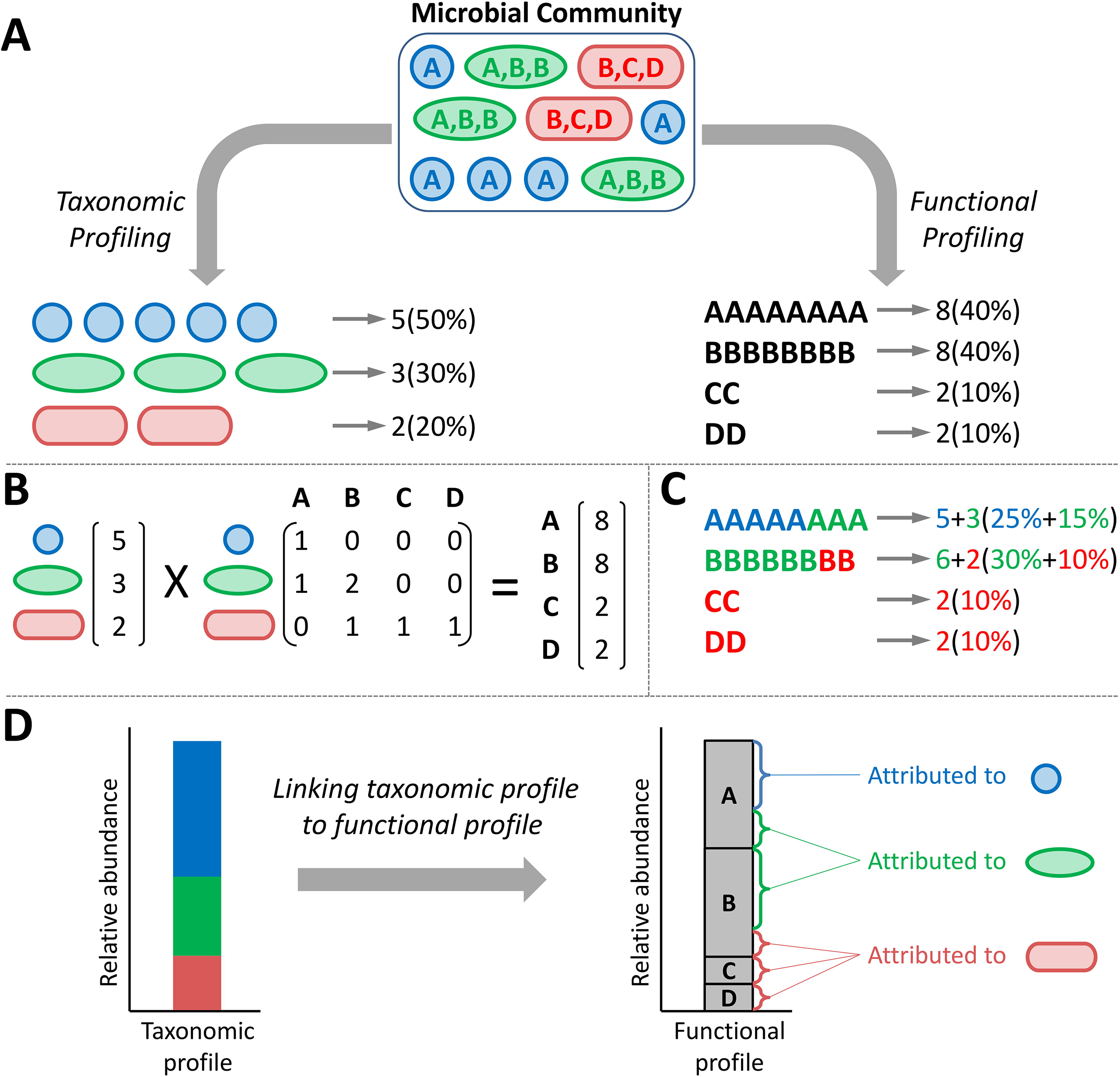
The taxonomic and functional compositions of a microbiome are inherently linked. **A)** Typical microbiome studies quantify and report the taxonomic (colored shapes) and functional (letters) profiles of a given community as separate entities. **B)** The functional profile of a community is a linear combination of the taxonomic composition and the genomic content of each taxon. **C)** Functional profiles can be deconvolved into taxon-specific functional profile, denoting which share of the abundance of each function is *attributed* to each taxon. **D)** Such deconvolved functional profiles can be visualized, illustrating the total abundance of each function as a stacked bar of taxon-specific attributions.

Importantly, however, recent years have witnessed an explosion of microbiome *multi-omic* studies that aim to describe simultaneously multiple aspects of community structure, including specifically both taxonomic and functional compositions (Greenblum et al., 2015; Huttenhower et al., 2012; Lloyd-Price et al., 2017; Pedersen et al., 2016; Taxis et al., 2015; Zhernakova et al., 2016). Moreover, recently developed methods can now determine both the taxonomic and functional profile of a given community from the same sequencing data, for example, by assigning shotgun metagenomic reads both taxonomic and functional annotations (Abubucker et al., 2012). Notably, these two facets of microbiome composition are not independent since the set of genes found in a metagenome and their abundances is a direct result of the set of genes (and their copy number) encoded by each community member and the relative abundance of each member in the community (Figure 1B). Put differently, the abundance of each gene family (or ‘function’) in the metagenome can be deconvolved into taxon-specific functional profiles in which shares of the gene family’s total abundance are attributed to specific taxa of origin (Carr, Shen-Orr, & Borenstein, 2013)(Figure 1C). This link between taxonomic and functional compositions can be used, for example, to predict functional abundances from 16s rRNA-based taxonomic profiles (Langille et al., 2013) or to identify taxonomic drivers of disease-associated functional shifts (Manor and Borenstein, 2017b). Importantly, this inherent relationship between taxonomic and functional profiles must also be considered when exploring how functional capacity co-varies with taxonomic composition across samples, since differences in gene abundances between samples are mainly derived from differences in taxonomic composition (Bäckhed et al., 2015; Oh et al., 2014; Turnbaugh et al., 2009). Yet, despite the growing appreciation for this link between taxonomic and functional compositions, an integrative tool that can simultaneously visualize both taxonomic and functional data and that can *account for* and *expose* the relationships between taxonomic and functional variation is lacking.

Here we introduce BURRITO (Browser Utility for Relating micRobiome Information on Taxonomy and functiOn), a web-based visualization tool that enables easy and intuitive exploratory analysis of the relationships between taxonomic and functional abundances across microbiome samples. BURRITO simultaneously provides a traditional interface for exploring taxonomic and functional abundances independently while also visualizing the links between these two microbiome facets and highlighting the share of each function’s total abundance that is attributed to each taxon (Figure 1D). Through an interactive interface, BURRITO also provides ample and precise information about such *attributions* (*i.e.*, the share of each function’s total abundance attributed to each taxon), as well as various summary statistics. To facilitate interactive data exploration and publication-quality figure generation for a wide audience, BURRITO further offers multiple options for data input and supports customizing various aspects of the visualization.

## Methods and Implementation

### User input and taxa-function mapping

BURRITO accommodates multiple types of input data and, depending on the provided data, uses different approaches to attribute the provided or inferred function abundances to taxa of origin (see Figure 1D). Specifically, the user can select one of three options for input data and for determining taxa-function attributions (Figure 2). In the first option, which requires the bare minimum in terms of input data, the user can simply provide a table of taxonomic abundances across samples (measured as either absolute read counts or relative abundances) using Greengenes 97% Operational Taxonomic Unit (OTU) IDs for each taxon (DeSantis et al., 2006). Given these taxonomic data, BURRITO will automatically predict the functional profile of each sample and will determine each function’s taxonomic attributions using a database of pre-annotated genomic content (following the approach described in Figure 1B-C). Briefly, in this approach, taxonomic abundances for each sample are first corrected by dividing each taxon’s abundance by its 16S rRNA copy number. The abundance of a given function that is attributed to each taxon (and ultimately the total abundance of that function) is then inferred by multiplying the corrected taxonomic abundances by the number of genes associated with that function in each taxon’s genomes. Data on the gene content in each taxon’s genome is obtained from PICRUSt (Langille et al., 2013b) and functional annotations are based on KEGG Orthology groups (Kanehisa et al., 2015; Kanehisa and Goto, 2000).

**Figure 2.**
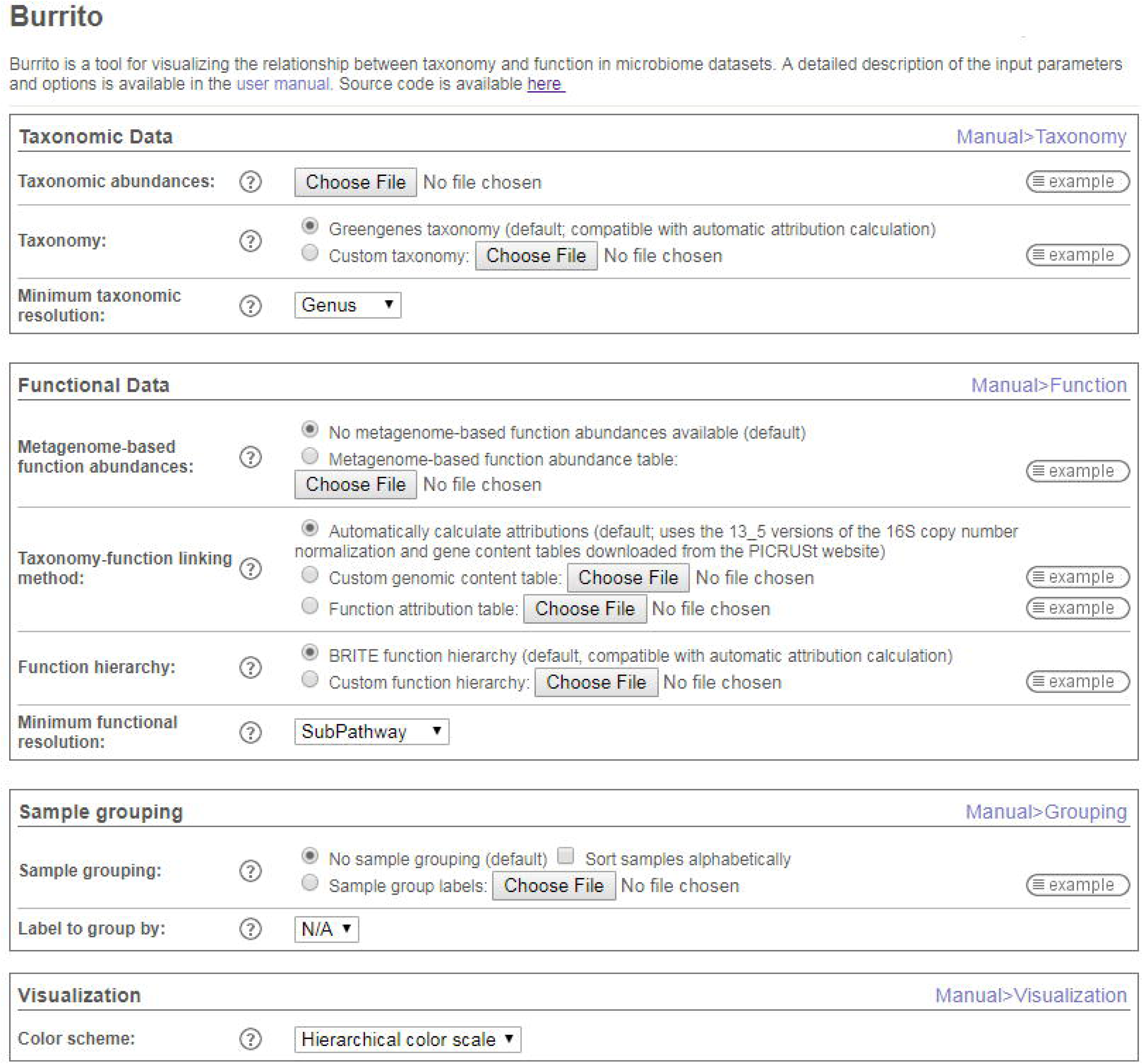
BURRITO’s upload page, describing the various input approaches BURRITO supports and other visualization and analysis options.

The second available option is similar to the one described above, but relies on user-provided genomic content annotations (instead of PICRUSt-based inferred content) to calculate functional profiles and the share of the functional profile attributed to each taxon. This approach is appropriate, for example, for exploring communities whose members may not be adequately represented by PICRUSt-inferred genomes but can be better characterized based on user proprietary data. As in the first option, the user is required to provide a taxonomic abundance table, but also provides a custom genomic content table describing the set of genes (and their copy number) encoded by each taxon. Given these data, functional abundances and attributions are calculated in the same manner as described above. When using this option, it is also assumed that taxonomic abundances are already corrected for 16S rRNA copy number. Moreover, using this approach the user is not limited to Greengenes OTU IDs or to KEGG Orthology groups, and alternately can use their taxonomic and/or functional classification of choice (as long as the same IDs are used in all relevant input files). Notably, when using custom taxonomic classification or custom hierarchical function relationships, the user can also provide files describing these custom systems to allow taxonomic and functional data to be grouped at different levels (see section ‘Exploring attributions at different taxonomic and functional levels’ below).

The third and final option relies on a pre-determined table of taxon-specific functional attributions rather than calculating attributions from taxonomic composition. This approach may be appropriate, for example, in cases where the user wishes to introduce specific custom modifications to a pre-calculated attribution table. Using this option requires a taxonomic abundance table (as above) and, instead of a genomic content table, a table of taxa-specific functional abundance attributions. As in the second approach, any taxonomic classification or functional hierarchy system can be used.

The specific format for each data input file and the specific restrictions associated with each approach are all noted on BURRITO’s upload page and are described in more detail in BURRITO’s documentation. Example data files are also available for download from the upload page. Notably, in all three approaches, the user can also upload an independent, paired dataset of functional abundance profiles (i.e. based on functional annotation of shotgun metagenomic sequencing), which will be visualized as described below alongside the calculated taxa-based functional profiles.

In addition to the above primary input, BURRITO’s upload page (Figure 2) further includes several visualization options, allowing users to better control the way data will be displayed. Specifically, BURRITO supports grouping samples based on user-provided labels (e.g., cases vs. control or conditions). Additionally, the user can select the minimum taxonomic and functional resolution to be displayed. Using a finer resolution allows exploring the data in more depth, but could slow visualization performance due to the large number of elements that need to be calculated, managed, and displayed. Finally, users can choose between hierarchical or random color schemes to distinguish taxa and functions.

### Visualizing the relationship between taxonomic and functional abundances

BURRITO uses two stacked bar plots, one for taxonomy (Figure 3A) and one for function (Figure 3B), to provide a standard visualization of taxonomic and functional relative abundances in each sample. Taxonomic abundances are taken from the user-provided taxonomic abundance table. Functional abundances are calculated as described in the previous sections and are displayed as the sums of all taxon-specific attributions for each function and sample. Precise abundance values for each taxon or function in each sample can be viewed in a *tooltip* that appears when the user hovers over any bar segment (Figure 3C). Additionally, hovering over a bar segment highlights the corresponding taxon’s or function’s bar segment across all samples to aid visual comparison of abundances between samples (Figure 3D).

**Figure 3.**
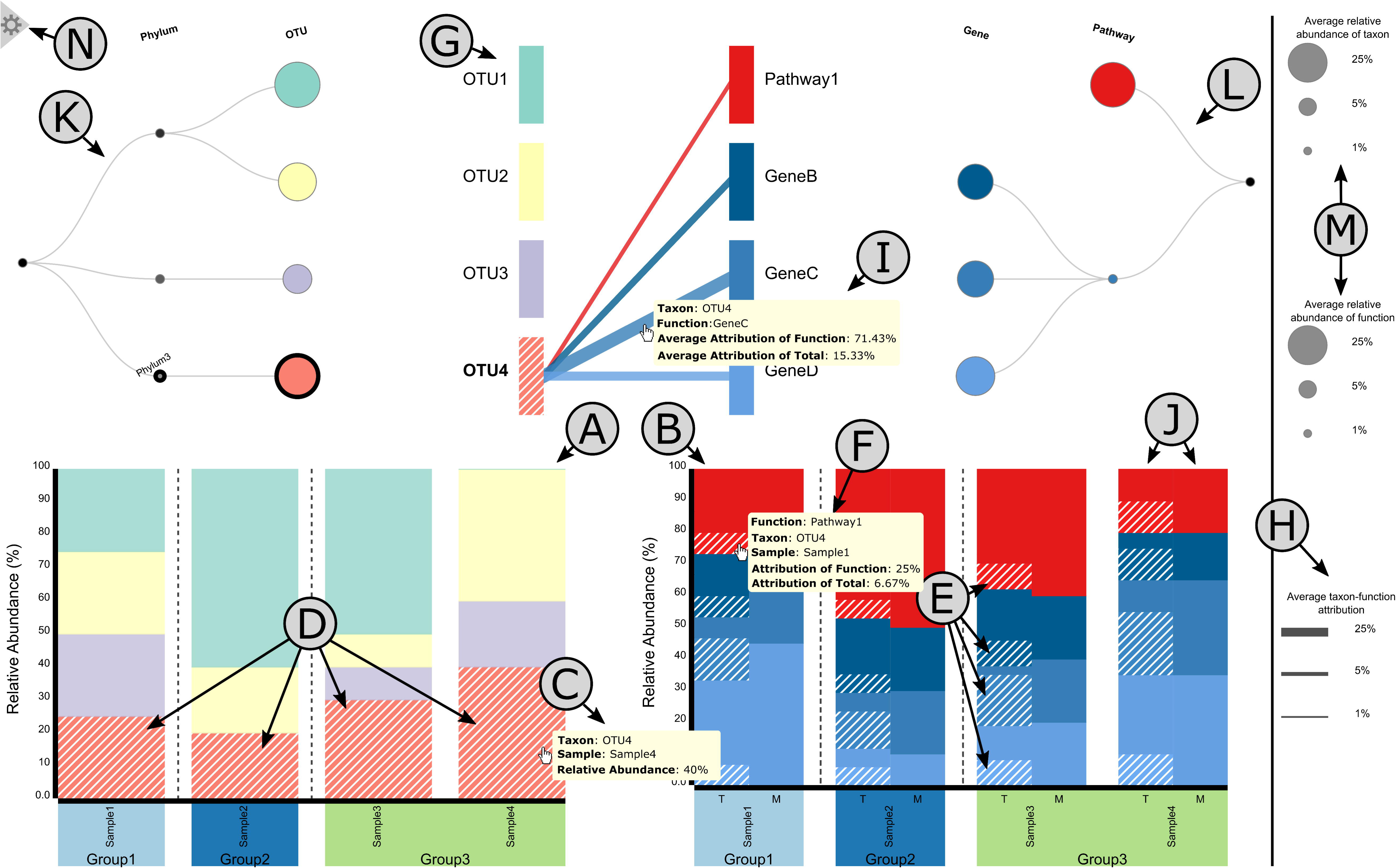
A layout of BURRITO’s visualization. **A, B)** Stacked bar plots of taxonomic and functional composition across samples. **C)** Tooltips appear when hovering over each taxon, providing information about this taxon’s relative abundance in each sample. **D)** Interactive highlighting of individual taxa, which correspondingly highlights the shares of functional abundances that can be attributed to the taxon in question **(E)**. **F)** Tooltips provide detailed function abundance and attribution data for each sample. **G)** The bipartite graph control panel identifies individual taxa and functions and shows links between them. **H)** Edge width in the control panel represents the *average* share of a function that is attributed to a given taxon. **I)** Exact taxon-function attribution values can be seen by hovering over the edge connecting the taxon to the function. **J)** An independent dataset of shotgun metagenomics-derived function abundances can be provided and displayed alongside inferred taxon-specific function abundances. **K, L)** The data can be viewed at a higher or lower taxonomic or functional resolution by clicking on nodes in the corresponding tree diagrams. **M)** The size of each node in the tree represents the average abundance of that entity. **N)** Opening the hidden menu allows users to export visualization plots and processed data.

The most innovative component of BURRITO is the visualization of how function abundance shares are attributed to the various taxa. This information is revealed when the user hovers over a bar segment in the taxonomic abundance bar plot, which, in addition to highlighting taxon abundances as noted above, also highlights the portion of each function abundance bar segment (in each sample) that is attributed to this taxon (Figure 3E). To view the exact function abundance share attributed to a given taxon, the user can click on (rather than hover over) a taxon’s bar segment to lock this taxon-specific attribution highlighting, and then hover over the highlighted portion of a function bar segment, revealing a tooltip with the corresponding information (Figure 3F).

BURRITO also displays a “control panel” that can be used to investigate specific taxa and/or functions and explore average abundances across samples (Figure 3G). Specifically, taxa and functions are represented by bars on the left and right sides of a bipartite graph. Hovering over the control panel performs similar highlighting as the bar plots, highlighting the linked abundances of individual taxa and functions and displaying (via a tooltip) the average relative abundance of each taxon or function. Moreover, when a taxon is highlighted, edges that link that taxon to all the functions it encodes are displayed, providing an easy reference for identifying the functions with shares attributed to that taxon. Similarly, when a function is highlighted, edges that indicate which taxa encode that function (and hence have shares of that function attributed to them) are displayed. The width of an edge between a taxon and a function represents the average share of that function’s abundance that is attributed to that taxon across all samples (Figure 3H). Clicking on a specific taxon or function in the control panel (or bar plots) locks the selection of that taxon or function, allowing the user to hover over each edge and view (via a tooltip) the exact taxon-function attribution values (Figure 3I). Additionally, when a taxon or function selection has been clicked (i.e., selected) and edges connecting taxa and functions are displayed, the user can highlight the abundances of a single function attributed to a single taxon by clicking on the edge between them.

BURRITO also supports comparison between taxa-based function abundances (calculated based on taxonomic profiles and genomic content as noted above) and a separate dataset of user-provided function abundances (typically obtained by functional annotation of shotgun metagenomic sequencing reads). If such a dataset is included in the input, these user-provided function abundances are displayed adjacent to the taxa-based function abundances for comparison (Figure 3J).

### Exploring attributions at different taxonomic and functional levels

BURRITO provides an interactive and intuitive interface for exploring abundance and attribution data at varying taxonomic and functional resolutions. Taxonomic classification and hierarchical function relationships (either those used by default or custom systems provided by the user as noted above) are each represented as a tree above the corresponding abundance bar plot (Figure 3K and 3L, respectively). To aid the user in understanding how different taxa or functions are related in the bipartite graph and bar plots, these trees are aligned to the left and right of the bipartite graph with all taxa and functions appearing in the same vertical order across all components of the visualization. These trees also indicate average taxon and function abundances across samples by the size of leaf nodes in the trees (Figure 3M). Beyond visualizing hierarchical relationships, these trees also provide a tool for interactive data exploration by allowing the user to expand or collapse different leaves or branches of each tree. Clicking on a leaf node reveals all taxonomic or functional subcategories of that node in the tree, and correspondingly expands the bipartite graph and subdivides the relevant relative abundance bars in the bar plot into the relative abundances of those subcategories. Alternatively, clicking on a non-leaf node within a tree performs the reverse operation, collapsing all visible descendants of that node, making the clicked node a leaf node, and aggregating the abundance bars for those descendants into the abundance bars for the clicked node. Importantly, these interactive features allow the user to dynamically drill up or down in both taxonomic and functional resolution across the different branches of either tree as they explore the data.

### Exporting visualization plots and processed data

BURRITO also provides options for exporting a static version of the visualization (e.g., for including in presentations or publications) and for downloading the function abundance table or attribution table underlying the displayed function plot. These options can be accessed from the visualization screen via a hidden menu (Figure 3N). Exported figures will maintain any currently-selected highlighting and taxonomic or functional tree expansion. In addition to exporting the full visualization, users can choose to individually export either bar plot of relative abundances and either half of the bipartite graph, which can serve as a legend for the color-coding of taxa or functions in the bar plots. All images can be exported in PNG or SVG format. If users wish to further explore the predicted function abundance or attribution data in more detail, they can also download the tables underlying the visualization. Function abundance and attribution tables can be downloaded at the minimum functional resolution (and minimum taxonomic resolution for the attribution table) specified on the upload page.

### Technical Implementation

BURRITO’s client is a browser page written in HTML and Javascript, utilizing the d3.js library to display data. User-submitted data are uploaded to an R Shiny server for processing (including, specifically, calculation of attributions) and the results are sent back to the browser for visualization. Additional details concerning BURRITO implementation can be found in the Supplementary Text.

## Case Studies & Discussion

To demonstrate the utility of BURRITO, we describe below its application to two microbiome datasets with varying properties.

### Case study 1: Exploring the effects of antibiotic treatment and recovery on inferred functional composition in the mouse cecum

We used BURRITO to visualize a publicly available dataset of 16S rRNA sequencing data, describing cecal samples from mice treated with antibiotics 2 days and 6 weeks after treatment (labeled ‘Abx Day 2’ and ‘Abx Day 42’, respectively), and time-matched controls (labeled ‘Control Day 2’ and ‘Control Day 42’, respectively) (Theriot et al., 2014). This study, which focused on associations of the microbiome with metabolomic data and colonization resistance, confirmed significant community perturbations in response to antibiotics. This dataset is also used in BURRITO’s *Preview* option on the upload page (see Figure 2), allowing users to examine BURRITO’s visualization and functionality (and to compare those to the examples provided in this case study) without the need to provide any additional data.

Given this dataset, we used the first input approach described above, allowing BURRITO to predict functional abundances (and taxon-specific attributions) based on the default PICRUSt-derived genomic content table. BURRITO’s visualization of taxonomic and functional profiles revealed relatively subtle functional variation despite drastic taxonomic variation across samples, a pattern commonly observed in microbiome studies (Manor and Borenstein, 2017a). Specifically, while Abx Day 2 samples are markedly different from, for example, Control Day 2 samples (with the former being dominated by species from the class *Bacilli* and the latter by species from the classes *Clostridia* and *Bacteroidia*), their predicted functional profiles are relatively similar (Figure 4A). Hovering over the various taxa in the taxonomic profiles highlighted the shares of the functional profile in the various sample groups that are attributed to microbes from these three classes (*Bacilli*, *Clostridia,* and *Bacteroidia*), demonstrating, for example, that attributions to the *Bacteroidia* species were relatively small compared to their abundance. See, for instance, sample NonAbx29, in which *Bacteroidia* is more abundant than *Clostridia* (54.93% vs. 44.13%), but has a smaller share attributed to it than to *Clostridia* in every functional category (Figure 4B). To further illustrate BURRITO’s functionality, we then visually searched for functions that are nonetheless differentially abundant between Abx Day 2 samples and other samples, focusing specifically on pathways in the metabolism category. We observed, for example, that Abx Day 2 samples were generally depleted for amino acid metabolism genes (Figure 4C). Selecting this pathway (by clicking on it in the bar plot or in the control panel) and examining the share of this pathway attributed to each taxon (by then clicking on the edge connecting this pathway to each taxon in the control panel), suggested that its depletion in Abx Day 2 samples could be explained by the fact that *Bacilli* contribute less than *Clostridia* to this pathway compared to their abundances. Specifically, we noted that the share of this pathway attributed to *Bacilli* in Abx Day 2 samples is smaller than the share of this pathway attributed to *Clostridia* in Abx Day 42 samples, even though the abundance of *Bacilli* and *Clostridia* in Abx Day 2 and in Abx Day 42 samples, respectively, is comparable (close to 100%) (Figure 4D). Similarly, the share of this pathway attributed to *Bacilli* in Abx Day 2 samples is comparable to the share of this pathway attributed to *Clostridia* in Control samples, even though the abundance of *Bacilli* in Abx Day 2 samples is higher than the abundance of *Clostridia* in control samples. This lower proportional contribution indicates a lower number of genes involved in this pathway in *Bacilli* compared to *Clostridia*. Lastly, we searched for additional functions with higher or lower share attributed to *Bacilli* by selecting this taxon (again, by hovering over or clicking on it in the control panel or in the bar plot) and examining the width of the attribution edges connecting it to each function. In this setting, it was easy to note that a relatively small share of the cell motility function is attributed to this taxon, compared to, for example, the shares of metabolic functions attributed to it (Figure 4E).

**Figure 4.**
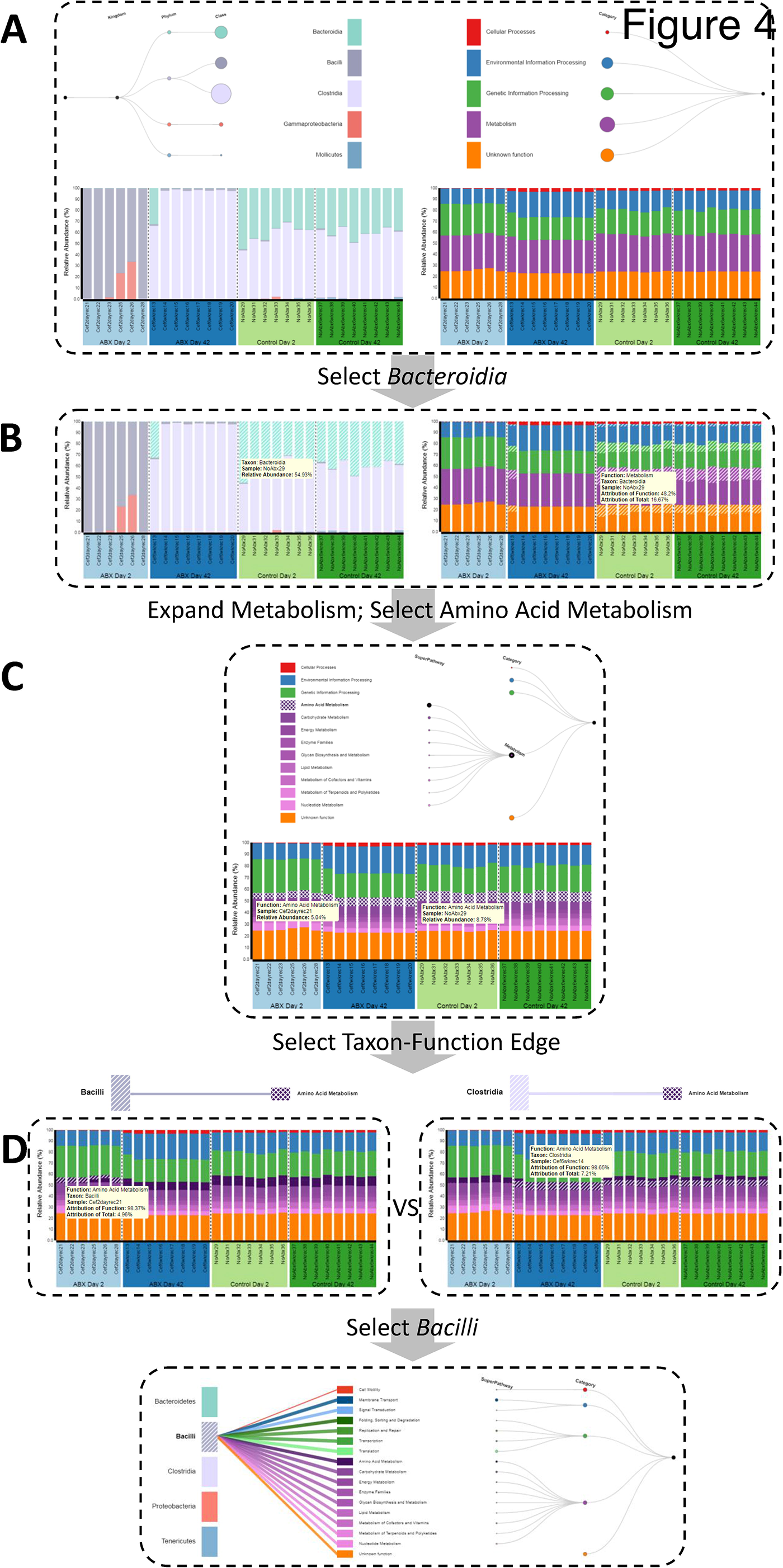
Using BURRITO to visualize the functional impacts of microbiome shifts in response to antibiotics. **A)** An overall view of BURRITO’s display for this dataset of 29 mouse cecal samples. **B)** Selecting a taxon (e.g., *Bacteroidia*) displays the share of each function attributed to this taxon. Tooltips provide exact attribution values. **C)** Expanding functional resolution and clicking on a given function (e.g., Amino Acid Metabolism) highlights this function in each sample. Tooltips provide exact abundance values. **D)** Once a function is selected, taxon-function edges in the control panel can be clicked, displaying the average share of the function attributed to the selected taxon and the sample-specific share of the function attributed to the selected taxon. **E)** Edges connecting a given taxon to all encoded functions provide information about the average share of each function attributed to this taxon.

### Case Study 2: Taxa-function relationships in the Human Microbiome Project

We additionally used BURRITO to visualize data from 21 supragingival plaque samples with both 16S rRNA and shotgun metagenomic data downloaded from the Human Microbiome Project (Huttenhower et al., 2012). We first used the 16S rRNA data alone (*i.e.*, again using the first input approach), examining inferred functional profiles and the taxa they are attributed to. As noted above, expanding the taxonomic tree can provide additional details about specific genera to which each function is attributed. Similarly, expanding the functional tree can offer insights into differentially abundant pathways and subpathways. For example, drilling down into subpathways in the Environmental Information Processing category and examining the average share from each subpathway attributed to the various phyla, we noted that the abundance of ABC transporter genes is attributed primarily to Actinobacteria, (average attribution across samples 24.87% of this function abundance), Firmicutes (24.65%), and Proteobacteria (29.55%). Indeed, samples with high relative abundance of these phyla tended to have higher abundance of this subpathway. Similarly, examining the shares of pyruvate metabolism (a subpathway of carbohydrate metabolism) attributed to genera from the phylum Proteobacteria revealed a specifically large attribution of the genus Neisseria, a prominent acid producer in the oral microbiome (Figure 5A). This is consistent with the capacity of strains from the genus Neisseria to metabolize lactate (via reactions included in the pyruvate metabolism pathway) (Hoshino and Araya, 1980; McLean et al., 2012).

**Figure 5.**
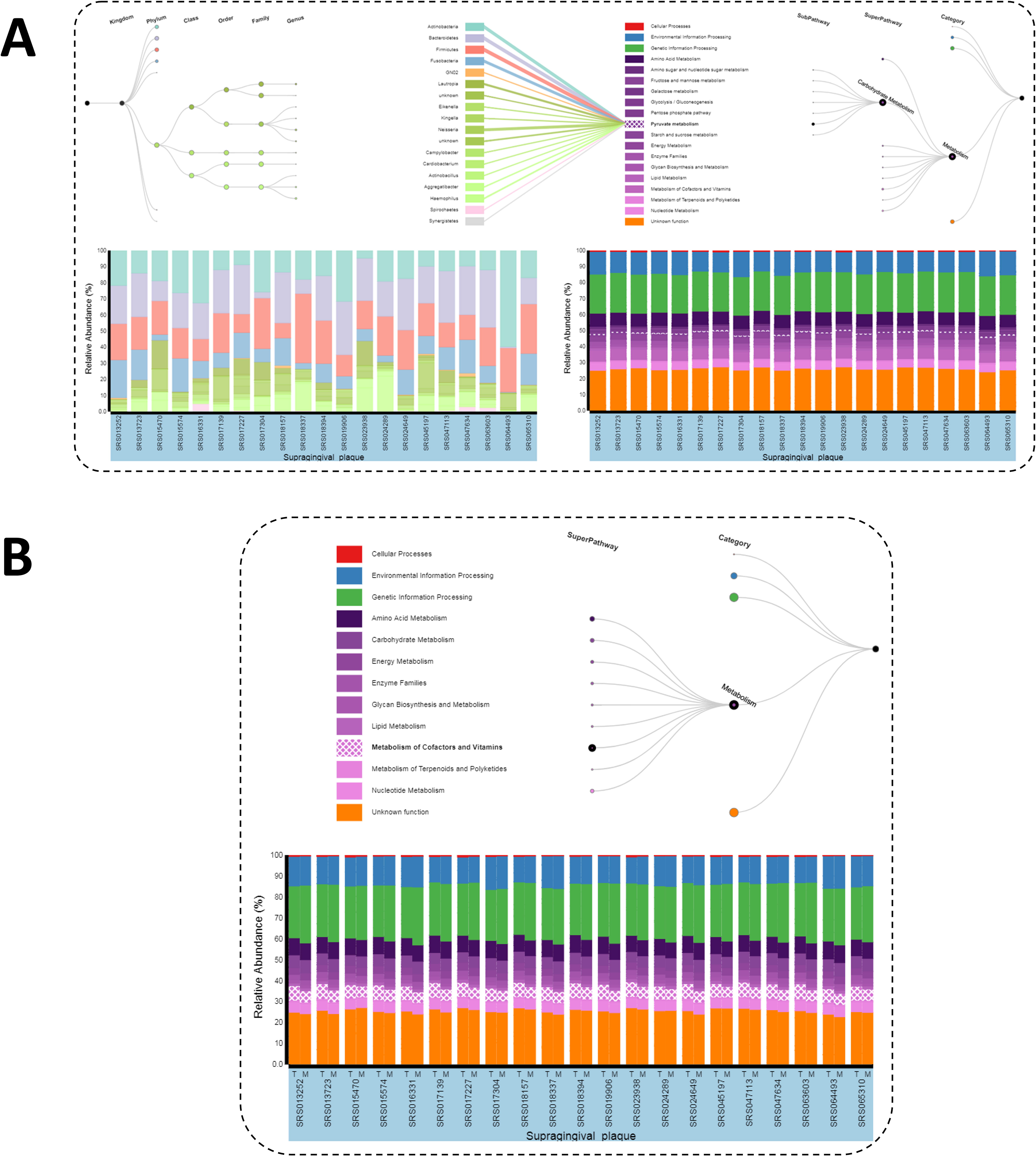
Taxa-function relationships across supragingival plaque samples from the Human Microbiome Project. **A)** Taxa and functions can be explored at different levels, using the trees in the control panel to expand taxa and functions of interest. This allows users to examine shares of specific functions that are attributed to specific taxa at high resolution. Edges in the control panel display information about the average share of a function attributable to each taxon across samples (with exact values provided via tooltips). **B)** Inferred taxa-based functional abundances (linked to 16S rRNA taxonomic data, T) can be displayed alongside measured functional relative abundance data obtained from metagenomic shotgun sequencing (M) for easy comparison.

Finally, Figure 5B demonstrates how BURRITO can also be used to compare such amplicon-based inferred functional profiles with functional abundance profiles obtained directly from shotgun metagenomic sequencing (when such functional profiles are provided as an additional input file). As expected, shotgun metagenomic-based profiles are generally in agreement with taxa-based inferred functional profiles, yet some differences can be observed, for example in the abundance of the genes related to the metabolism of cofactors and vitamins (highlighted).

## Conclusions

BURRITO is a web-based tool that addresses a major gap in currently available visualization tools for microbial ecology research. Studies in this field typically analyze, explore, and visualize taxonomic and functional abundances separately and fail to account for the interdependence between the two, potentially due to a shortage of tools that support simultaneous and integrative study of taxonomic and functional profiles. Our tool enables data exploration and hypothesis generation based on the attribution of functional abundances to specific taxa, providing a novel view into microbiome variation and dynamics.

## Availability and requirements

- **Project name:** BURRITO
- **Project home page:** http://elbo.gs.washington.edu/software_burrito.html
- **Operating system:** Platform independent
- **Programming language:** HTML 5, JavaScript, D3
- **Other requirements:** None
- **Compatible browsers:** Chrome version 61.0+, Firefox version 53.0+, Opera version 48.0+, Safari 11.0+ (PNG exports not supported in Safari)
- **License:** Non-commercial research use only (available at https://github.com/borenstein-lab/Burrito)

## List of abbreviations

BURRITO: Browser Utility for Relating micRobiome Information on Taxonomy and functiOn
OTU: Operational Taxonomic Unit
KEGG: Kyoto Encyclopedia of Genes and Genomes

## Authors’ contributions

CPM, AE, CN, and EB conceived and designed this study; CPM, AE, CN, and WGM implemented a preliminary version of BURRITO; CPM, AE, and CN completed the implementation of BURRITO; CPM, AE, CN, and EB wrote the paper. All authors read and approved this manuscript.

## Acknowledgements

A preliminary version of BURRITO was implemented as an assignment in a Data Visualization course instructed by Jeffrey Heer and we thank him and the course TAs for their feedback. We also thank all members of the Borenstein lab for helpful discussions and software testing, and Casey Theriot for the data displayed in one of the case studies.

## References

Abubucker, S., Segata, N., Goll, J., Schubert, A. M., Izard, J., Cantarel, B. L., et al. (2012). Metabolic reconstruction for metagenomic data and its application to the human microbiome. PLoS Comput. Biol. 8, e1002358. doi:10.1371/journal.pcbi.1002358.

Bäckhed, F., Roswall, J., Peng, Y., Feng, Q., Jia, H., Kovatcheva-Datchary, P., et al. (2015). Dynamics and Stabilization of the Human Gut Microbiome during the First Year of Life. Cell Host Microbe 17, 690–703. doi:10.1016/j.chom.2015.04.004.

Brewer, C. A. (2017). ColorBrewer: A web tool for selecting colors for maps. Available at: colorbrewer2.org [Accessed October 9, 2017].

Carr, R., Shen-Orr, S. S., and Borenstein, E. (2013). Reconstructing the genomic content of microbiome taxa through shotgun metagenomic deconvolution. PLoS Comput. Biol. 9, e1003292. doi:10.1371/journal.pcbi.1003292.

DeSantis, T. Z., Hugenholtz, P., Larsen, N., Rojas, M., Brodie, E. L., Keller, K., et al. (2006). Greengenes, a chimera-checked 16S rRNA gene database and workbench compatible with ARB. Appl. Environ. Microbiol. 72, 5069–72. doi:10.1128/AEM.03006-05.

Greenblum, S., Carr, R., and Borenstein, E. (2015). Extensive Strain-Level Copy-Number Variation across Human Gut Microbiome Species. Cell 160, 583–594. doi:10.1016/j.cell.2014.12.038.

Hoshino, E., and Araya, A. (1980). Lactate degradation by polysaccharide-producing Neisseria isolated from human dental plaque. Arch. Oral Biol. 25, 211–212. doi:10.1016/0003-9969(80)90023-0.

Huttenhower, C., Gevers, D., Knight, R., Abubucker, S., Badger, J. H., Chinwalla, A. T., et al. (2012). Structure, function and diversity of the healthy human microbiome. Nature 486, 207–214. doi:10.1038/nature11234.

Kanehisa, M., and Goto, S. (2000). KEGG: Kyoto Encyclopedia of Genes and Genomes. Nucleic Acids Res. 28, 27–30. doi:10.1093/nar/28.1.27.

Kanehisa, M., Sato, Y., Kawashima, M., Furumichi, M., and Tanabe, M. (2015). KEGG as a reference resource for gene and protein annotation. Nucleic Acids Res 44, D457–D462. doi:10.1093/nar/gkv1070.

Kuntal, B. K., Ghosh, T. S., and Mande, S. S. (2013). Community-Analyzer: A platform for visualizing and comparing microbial community structure across microbiomes. Genomics 102, 409–418. doi:10.1016/j.ygeno.2013.08.004.

Langille, M. G. I., Zaneveld, J., Caporaso, J. G., McDonald, D., Knights, D., Reyes, J. A., et al. (2013a). Predictive functional profiling of microbial communities using 16S rRNA marker gene sequences. Nat. Biotechnol. 31, 814–21. doi:10.1038/nbt.2676.

Langille, M. G. I., Zaneveld, J., Caporaso, J. G., McDonald, D., Knights, D., Reyes, J. A., et al. (2013b). Predictive functional profiling of microbial communities using 16S rRNA marker gene sequences. Nat. Biotechnol. 31, 814–21. doi:10.1038/nbt.2676.

Lloyd-Price, J., Mahurkar, A., Rahnavard, G., Crabtree, J., Orvis, J., Hall, A. B., et al. (2017). Strains, functions and dynamics in the expanded Human Microbiome Project. Nature. doi:10.1038/nature23889.

Manor, O., and Borenstein, E. (2017a). Revised computational metagenomic processing uncovers hidden and biologically meaningful functional variation in the human microbiome. Microbiome 5. doi:10.1186/s40168-017-0231-4.

Manor, O., and Borenstein, E. (2017b). Systematic Characterization and Analysis of the Taxonomic Drivers of Functional Shifts in the Human Microbiome. Cell Host Microbe. doi:10.1016/j.chom.2016.12.014.

McLean, J. S., Fansler, S. J., Majors, P. D., McAteer, K., Allen, L. Z., Shirtliff, M. E., et al. (2012). Identifying Low pH Active and Lactate-Utilizing Taxa within Oral Microbiome Communities from Healthy Children Using Stable Isotope Probing Techniques. PLoS ONE 7, e32219. doi:10.1371/journal.pone.0032219.

Oh, J., Byrd, A. L., Deming, C., Conlan, S., Barnabas, B., Blakesley, R., et al. (2014). Biogeography and individuality shape function in the human skin metagenome. Nature 514, 59–64. doi:10.1038/nature13786.

Ondov, B. D., Bergman, N. H., and Phillippy, A. M. (2011). Interactive metagenomic visualization in a Web browser. BMC Bioinformatics 12, 385. doi:10.1186/1471-2105-12-385.

Pedersen, H. K., Gudmundsdottir, V., Nielsen, H. B., Hyotylainen, T., Nielsen, T., Jensen, B. A. H., et al. (2016). Human gut microbes impact host serum metabolome and insulin sensitivity. Nature advance online publication. doi:10.1038/nature18646.

Robertson, C. E., Harris, J. K., Wagner, B. D., Granger, D., Browne, K., Tatem, B., et al. (2013). Explicet: graphical user interface software for metadata-driven management, analysis and visualization of microbiome data. Bioinformatics 29, 3100–3101. doi:10.1093/bioinformatics/btt526.

Taxis, T. M., Wolff, S., Gregg, S. J., Minton, N. O., Zhang, C., Dai, J., et al. (2015). The players may change but the game remains: network analyses of ruminal microbiomes suggest taxonomic differences mask functional similarity. Nucleic Acids Res., gkv973. doi:10.1093/nar/gkv973.

Theriot, C. M., Koenigsknecht, M. J., Carlson Jr, P. E., Hatton, G. E., Nelson, A. M., Li, B., et al. (2014). Antibiotic-induced shifts in the mouse gut microbiome and metabolome increase susceptibility to Clostridium difficile infection. Nat. Commun. 5. doi:10.1038/ncomms4114.

Turnbaugh, P. J., Hamady, M., Yatsunenko, T., Cantarel, B. L., Duncan, A., Ley, R. E., et al. (2009). A core gut microbiome in obese and lean twins. Nature 457, 480–484. doi:10.1038/nature07540.

Vázquez-Baeza, Y., Pirrung, M., Gonzalez, A., and Knight, R. (2013). EMPeror: a tool for visualizing high-throughput microbial community data. GigaScience 2, 16. doi:10.1186/2047-217X-2-16.

Wang, Y., Xu, L., Gu, Y. Q., and Coleman-Derr, D. (2016). MetaCoMET: a web platform for discovery and visualization of the core microbiome. Bioinformatics 2, btw507. doi:10.1093/bioinformatics/btw507.

Zhernakova, A., Kurilshikov, A., Bonder, M. J., Tigchelaar, E. F., Schirmer, M., Vatanen, T., et al. (2016). Population-based metagenomics analysis reveals markers for gut microbiome composition and diversity. Science 352, 565–569. doi:10.1126/science.aad3369.

